# Neural Basis of Odometry in Drosophila

**DOI:** 10.1101/2025.03.23.644782

**Authors:** Ilaria D’Atri, Shamik DasGupta

## Abstract

Path integration is a mode of navigation in which travel distance and direction are integrated to calculate position. Estimating travel distance, or ‘odometry,’ requires the summation of translational vectors over time. Although neurons sensitive to translational velocity have been identified in numerous species, a definitive relationship between the activity of these translational velocity neurons and the subjective perception of distance has yet to be established. We developed a new memory-based assay to dissect the neural circuit of distance estimation. Flies can estimate distance by using self-motion, associate it with aversive stimulus onset, and use the PFN→hΔB pathway to estimate it. Thus, we provide evidence for self-motion integration for distance estimation in flies and point to the neural basis of this integration.

## Main Text

During path integration, animals integrate self-generated translational and rotational vectors to calculate distances and determine travel directions **(*1*–*4*)**. While head-direction systems in vertebrates and insects have revealed how the brain integrates rotational vectors to calculate heading **(*5*–*8*)**, the neural mechanisms underlying distance estimation are not clearly understood **(*9*–*24*)**. While homing insects can estimate distance over impressive distances **(*25*– *29*)**, the extent to which non-homing insects, like the fruit fly *D. melanogaster*, can calculate distance remains unclear. To interrogate fruit flies’ ability to estimate distance during walking, we developed an associative condition assay (Fig. 1A, left) that required them to learn to associate distances with electric shock. We introduced single flies in a narrow annular arena where the continuity of the track was disrupted by a wall that spans across the track. The arena was kept inside a temperature-controlled incubator and was illuminated only with infrared LEDs to stop the flies from using visual inputs. We tracked the fly’s position in the chamber with machine-vision and delivered brief pulses of electric shock to the fly when it entered a pre-assigned portion (always displayed as the bottom left quarter during post-processing) of the chamber (Fig. 1A, right). Flies paced around the arena but, after training, began to avoid the shock-associated quarter of the chamber. Moreover, the distance they traversed before altering their course—hereafter referred to as “run lengths”—became notably shorter after training, compared to the distances they covered when subjected to shocks (Fig. 1B). To quantify these changes, we used two parameters: PI-Place measured the flies’ avoidance of the shock-associated part of the chamber, and PI-Distance quantified their tendency to terminate a walking bout before traversing shock-associated distances (Fig. 1C to E). Both indices were normalized by total time (PI-Place) or run number (PI-Distance) to scale them between -1 and 1. We found that run lengths during testing were tightly correlated with shock-associated run lengths, demonstrating that flies can not only estimate the distance they walk between successive changes in directions but also adjust this length under learning. While files showed distance and place learnings across a range of chamber sizes and shapes (Fig. 1B and S1A), their performances began to suffer when the track size reached ∼150 body lengths (340mm), indicating a limitation in the fly’s ability to path integrate over large distances or associate long distances and large patches with an aversive stimulus. We then tracked the decay kinetics of these memories (Fig. 1F). We observed that the place memory decays much faster while the distance memory stays stable for at least an hour, like reports of distance memory during walking from other insects **(*30, 31*)**. We hypothesize that these place and distance learning systems could exist independently, and perhaps place avoidance stems from locally deposited olfactory cues. This idea was further supported by the fact that only the place learning is affected when flies are displaced by relocating the wall after training (Fig. S1B) or when they make two full rotations in a wall-less chamber (Fig. S1C). We observed that largely anosmic mutants **(*32*)**, *orco*^*1*^, were deficient in place avoidance but showed normal distance memory, pointing towards a role of olfaction for place memory (Fig. 1G) **(*33*)**.

**Fig. 1.**
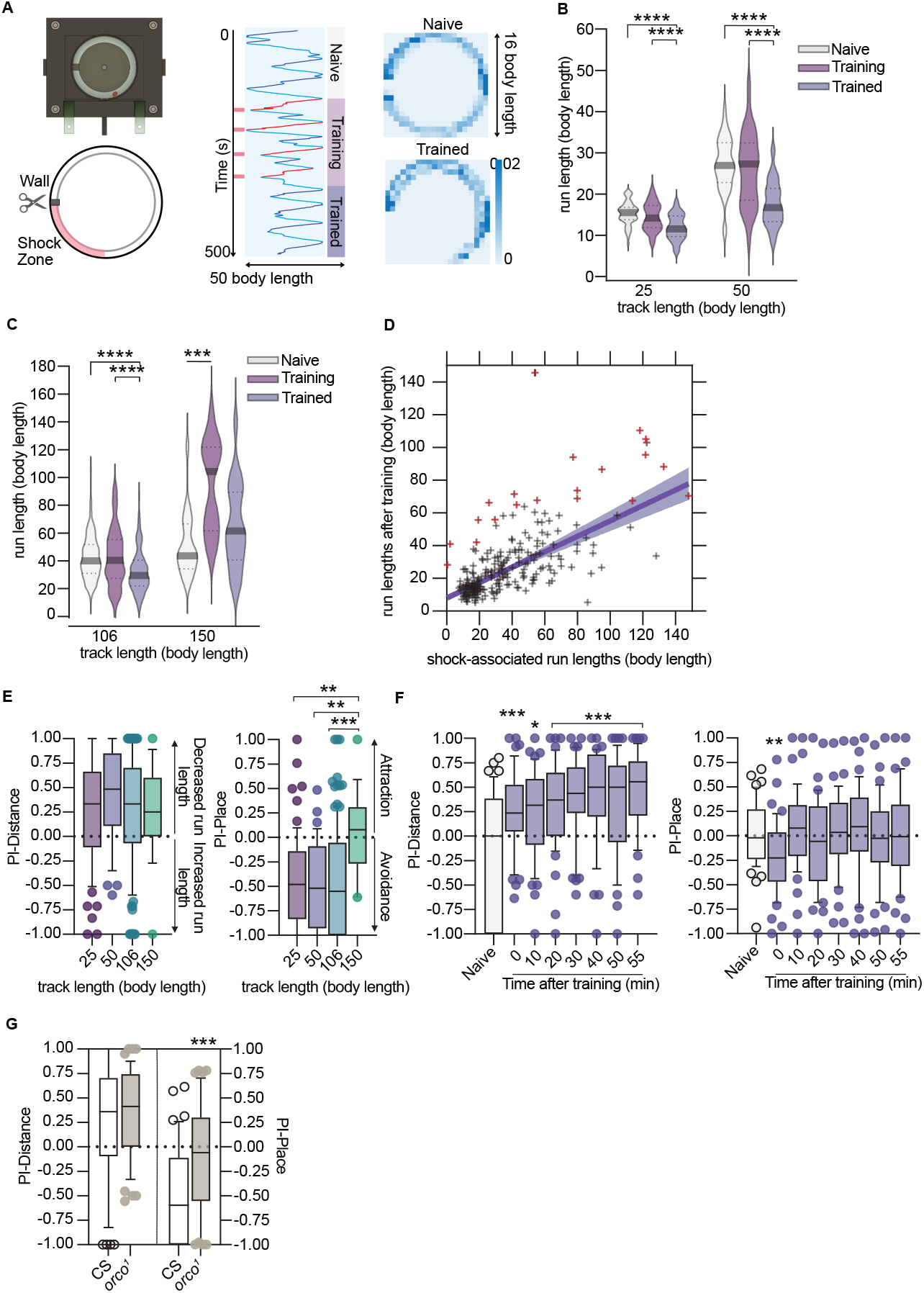
Place and distance learning in flies. (**A**) (Left) Top view of the 40cm diameter sized chamber: a movable wall breaks the continuity of the track and stops the fly from walking in full circles. The track length of this chamber is ∼50 body lengths. The red oval represents a fly drawn to scale. (Middle) Kymograph of a single fly; the tracks are linearized by cutting the circle at the wall position (scissor location). Individual runs are displayed in alternating shades of light and dark blue. Shock-associated runs are shown in red. During training, the fly receives electric shocks (pink ticks) if it resides in the bottom-left quarter of the chamber. (Right) Occupancy histogram of the same fly before and after training. The fly avoids the shock-associated bottom-left quarter of the chamber after training. (**B, C**) Flies walk shorter distances between successive turns after training. Asterisks indicate statistically significant differences from corresponding Naïve data. (**D**) Mean training and trained run length data from (B, C) were re-plotted pairwise to demonstrate a linear relationship between mean shock-associated run lengths and mean run lengths after training. The shaded region represents a 95% confidence interval of the fit. Red plus signs represent outliers. R^2^ = 0.39. (**E**) A positive PI-Distance reflects the shortening of the run lengths after training, and a negative PI-Place captures the avoidance of the shock-associated part of the chamber. Asterisks represent a significant difference from other groups. (**F**) One-hour time-course to follow the Distance (left) and the Place memory (right) decay. Place avoidance decays faster than distance memory. Asterisks represent a significant difference from naïve. (**G**) *orco*^*1*^ mutants are deficient in place avoidance but show normal distance learning. The asterisk represents a significant difference from the wild-type control. In all box and whiskers plots, the box extends from 25-75th percentiles with a median line, and whiskers extend from 10-90% percentiles. Data outside these ranges are displayed as individual points. Asterisks indicate significant differences using the convention * *p*<0.05, ** *p*<0.01, *** *p*<0.001, **** *p*<0.0001.

Having established the foundations of the assay, we sought to elucidate the neural circuit responsible for distance estimation. Numerous neurons in the fly brain respond to forward movement, making it challenging to determine which velocity-responsive neuron outputs are integrated for distance estimation. Thus, we focused on two brain regions implicated in associative learning **(*34*–*36*)** and movement encoding **(*37*–*41*)**: the mushroom bodies and the velocity-responsive cells of the central complex.

We employed the optogenetic inhibitor GtACR **(*42*)**, expressing it in targeted neurons and selectively silencing them during training. We first focused on the intrinsic neurons of the mushroom bodies, a well-characterized center for associative learning in flies. Inhibiting different subsets of Kenyon cells of the mushroom body (Fig. 2A) did not affect either place avoidance or distance learning. This finding indicates that, unlike olfactory aversive learning, the formation of aversive place avoidance is independent of mushroom body activity and potentially reliant only on local olfactory cues.

**Fig. 2.**
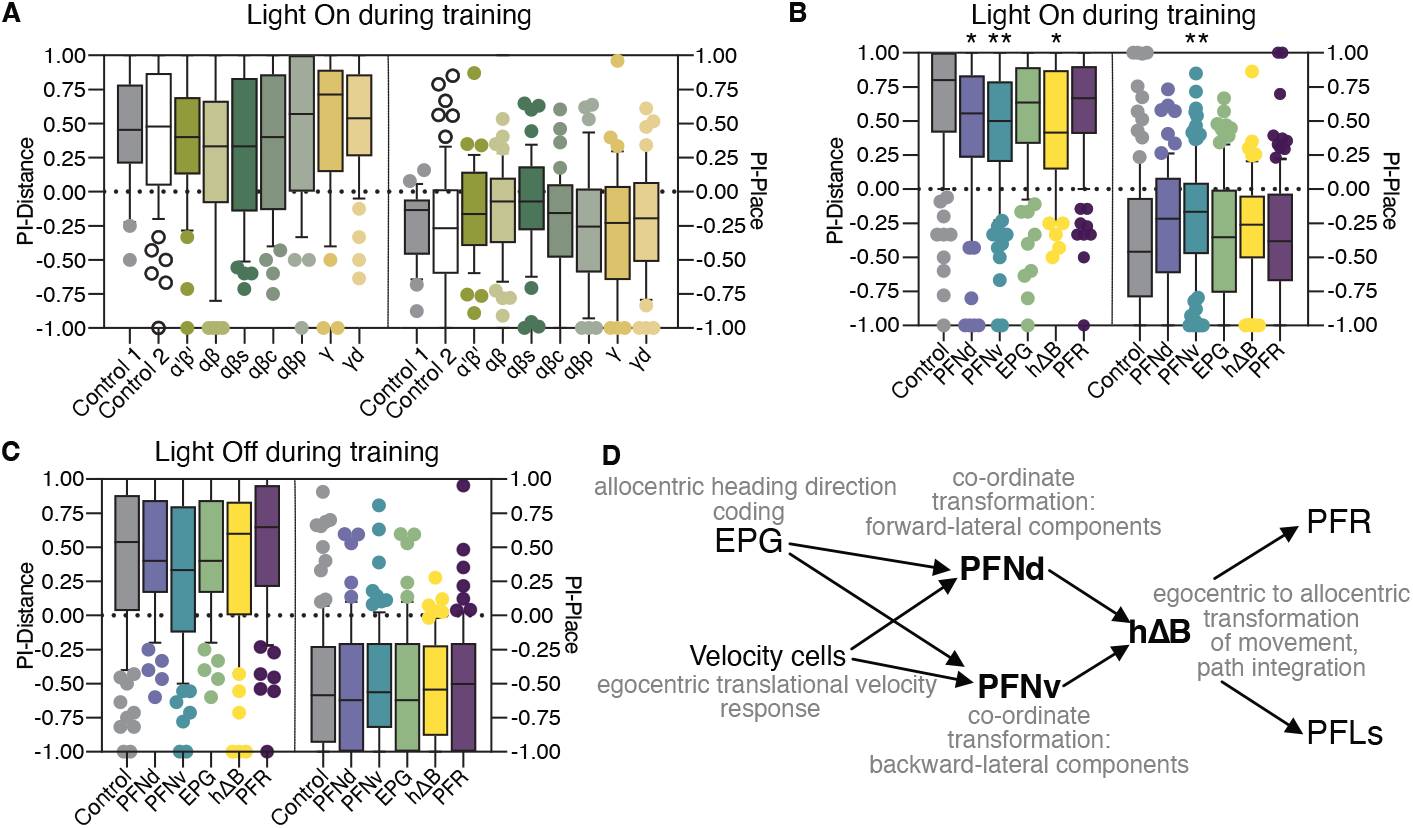
Neural circuit of distance memory acquisition. **(A)** GtACR-mediated optogenetic inhibition of the mushroom body neurons has no effect on place and distance learning. (**B**) However, similar manipulation of the central complex neurons affects distance memory acquisition. Asterisks denote statistically significant differences from the control. (**C**) Light-off controls for the optogenetic experiment in Fig 2B. **(D)** Simplified connectivity diagram and their proposed roles from past literature for the tested central complex neurons. Neurons that give phenotype in this assay are shown in bold. Box plot and Asterisks conventions are the same as in Fig 1.

Next, we investigated a subset of the central complex neurons, the PFN→hΔB pathway (Fig. 2B), for their proposed roles in navigation **(*15, 38*–*40*)**. Past works are consistent with the idea that the activity of the EPG neurons represents flies’ allocentric heading direction, and the PFN neurons represent flies’ translational vectors **(*15, 38*–*40*)**. PFNs have two distinct cell types: forward-ipsilateral velocity encoding PFNds and backward-contralateral velocity encoding PFNvs. In addition to translational velocity inputs, PFNs also receive inputs from EPG so that the egocentric translational velocities are referenced to an allocentric coordinate system **(*15, 38*– *40*)**. The hΔBs combine the PFNds and PFNvs outputs to represent the net translational velocity of the fly. Previous work demonstrated that chronic disruption of PFNds’ activity interferes with food-search behavior in the dark, indicating a role for PFNds in path integration **(*38*)**. PFR neurons principally receive inputs from hΔB cells, but their role in fly behaviors remains unclear **(*8, 38, 39*)**. Blocking outputs of the translational velocity-responsive neurons of the PFN→hΔB pathway (Fig. 2C and D) disrupted the formation of accurate distance memory, leaving place memory largely intact. We, therefore, hypothesized that the outputs of the hΔBs are integrated to compute the distance during shock association.

If hΔB inputs are integrated to calculate the shock-associated distance, transiently increasing hΔB activity via optogenetics should make the perceived shock-associated distance longer. This, in turn, would increase the chances of producing run lengths longer than the actual shock-associated run length during testing and worsen the Performance-Distance. We expressed the optogenetic activator CsChrismon (CsChR) in hΔB neurons and focused our activating light in different positions along the arena. Transient current injection into hΔB neurons near the shock-associated zone (Fig. 3A) increased error in distance memory, supporting our idea that their output is integrated to calculate the shock-associated distance.

**Fig. 3.**
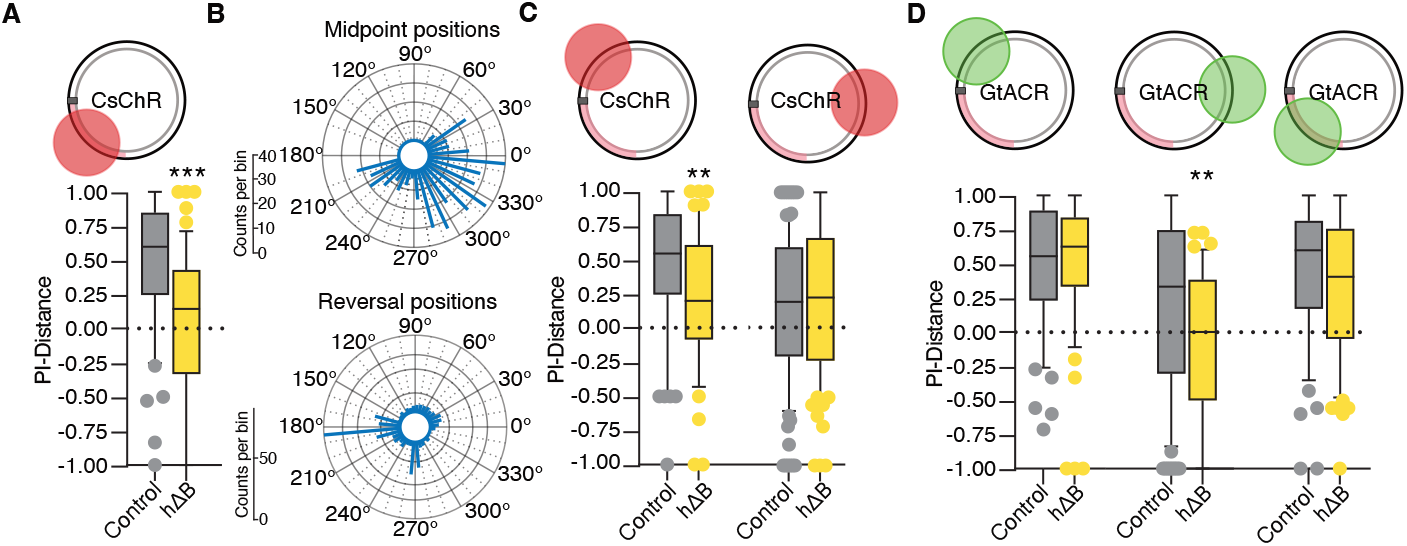
hΔB activity determines the estimated distance during memory acquisition. (**A**) Transient current injection into hΔB during training worsens performance during testing. Asterisks indicate a significant difference from the control. (**B**) Distribution of run midpoints and reversal positions during training. Data from the control flies from Fig 2C is analyzed for the positions. (**C**) During training, optogenetic current injection into the hΔB neurons near reversal points worsens performances during testing (2A, C-Left). Asterisks indicate a significant difference from the control. (**D**) However, optogenetic inhibition only affects performance when the light is focused near run midpoints. Box plot and Asterisks conventions are the same as in Fig 1.

This approach also allowed us to test whether hΔB outputs are uniformly integrated during walking. Since reversals are non-uniformly distributed along the chamber (Fig. 3B), we chose our stimulation zones to overlap with either the reversal points or the midpoints of the runs. Activating hΔB neurons near the reversal points of the runs away from the shock-associate zone increased error in distance memory (Fig. 3A, C). In contrast, optical current injection of hΔB neurons in the middle of the chamber (Fig. 3D) resulted in no change in distance memory. Interestingly, results from transient inactivation of hΔB followed the opposite pattern. Inhibition near the midpoints produced a significant drop in the distance performance index, whereas inhibition near the start/end points had marginal to no effect on PI-Distance. Together, these results indicate that the hΔB integration is non-uniform, and as suggested by prior work, the path integrator is zeroed as the fly nears the run midpoint **(*17*)**.

Next, we focused on the role of the PFN→hΔB pathway in distance memory retrieval. Like the previous experiment, we used the optical current injection approach to inject additional activity into these neurons. Since runs were unevenly distributed during testing (Fig. 4A), we categorized each run based on the distance of the center of the optogenetic stimulation from the run’s starting position (Fig. 4B). Transient current injection to the hΔB neurons reduced run lengths when the light is focused near reversal points, but run lengths are increased when hΔB>ChR expressing flies receive optogenetic stimulations near run midpoints (Fig. 4C, E). These data are consistent with hΔBs’ role in odometry and support the model that the odometer is zeroed near run midpoints. Interestingly, in naïve flies, transient current injection decreases turn-to-turn distances in all stimulation locations (Fig. 4D, E), pointing toward at least two possibilities. Either uniform odometry between turns occurs in naïve flies where the odometer is zeroed at the reversal points, or increased hΔB activity directly increases reversal probability in naïve flies.

**Fig. 4.**
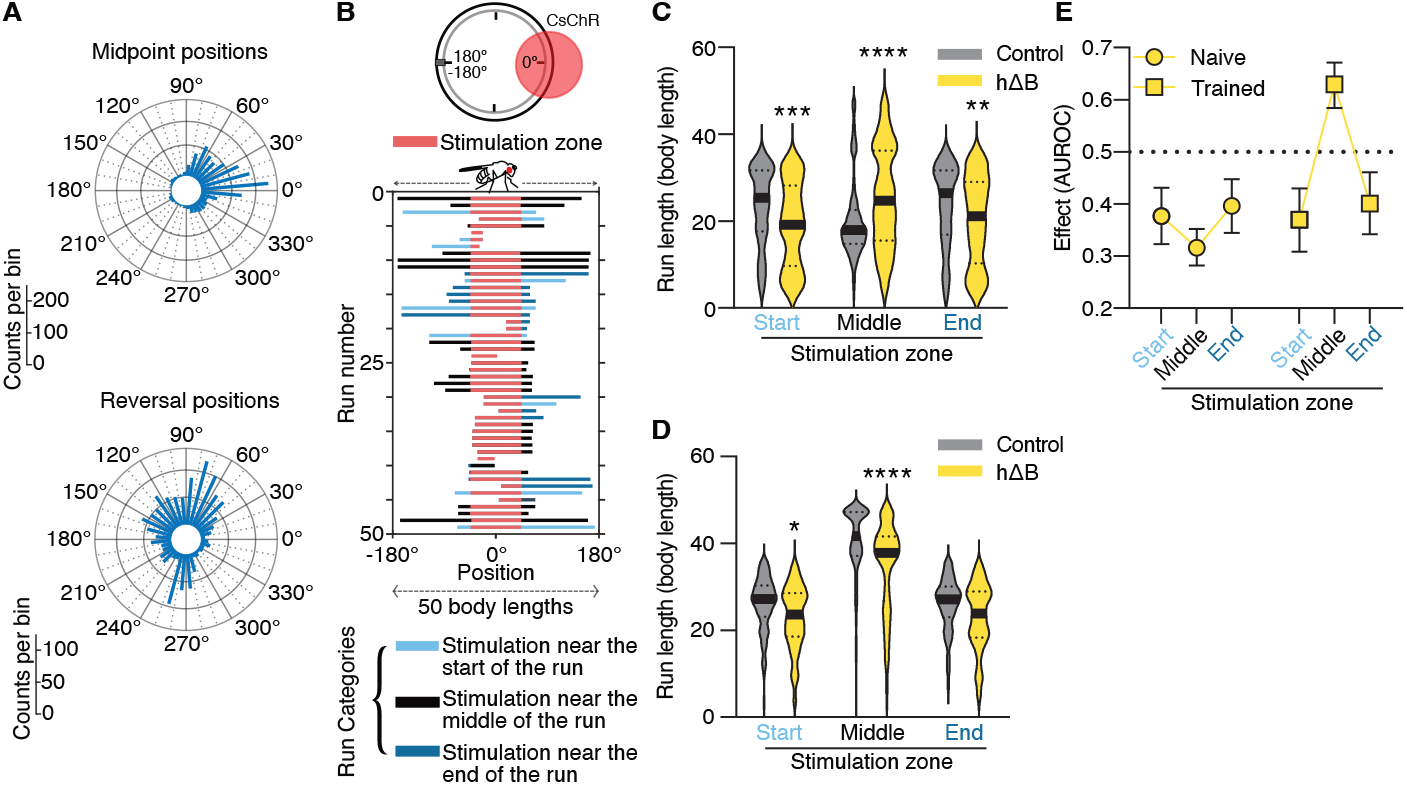
hΔBs have a role in memory retrieval. (**A**) Distribution of run midpoints and reversal positions during testing. Data from the control flies from Fig 2C is analyzed for the positions. (**B**) Experiment outline and example linearized runs from the control flies after training. The runs are binned on the relative distance of the stimulation zone’s midpoint relative to the run’s starting point. (**C, E**) In trained files, optogenetic stimulation of hΔB neurons after distance learning causes a decrease in run length when the stimulation zone is located near reversal locations. However, run length is increased when hΔB neurons are stimulated near run midpoints. **(D, E)** In naïve flies, optogenetic stimulations of hΔB neurons reduce run lengths, regardless of the stimulation zone position. Asterisks indicate significant differences relative to the matched control: * *p*<0.05, ** *p*<0.01, *** *p*<0.001, **** *p*<0.0001.

We demonstrate that *D. melanogaster* can estimate distances during walking and can learn to change travel distances after associative conditioning. We show that velocity-responsive neurons of the PFN→hΔB pathway are critical for distance estimations, and output hΔB neurons are integrated for distance estimation. This circuit also has a role in terminating walking bouts to generate turns in both naïve and trained flies, but through different mechanisms. Our data captures several key features of previous models of path integration and provides a mechanistic link. We speculate that flies employ separate neural pathways to store and retrieve path integration vectors and mechanism(s) for zeroing the path integrator changes after associative learning.

## Acknowledgements

We thank Harry Mayes, William D. Constance, Hoi Lam Wong, and Michelle Vermeulen for their help with data acquisition. We also thank Sadra Sadeh for his comments on the manuscript and Gero Miesenböck, Helen Weavers, and Scott Waddell for sharing fly lines.

## Data and materials availability

All data from the figures and analysis codes will be available through Zenodo and Github upon peer-reviewed publication. The corresponding author will make other materials available upon request.

## Supplementary Materials

Materials and Methods

Figs. S1

Tables S1 to S2

