## Supplementary Material for "Neural Basis of Odometry in Drosophila"

**The PDF file includes:**

Materials and Methods
Fig. S1
Tables S1 to S2

### Materials and Methods

Fly husbandry and strains

All the experiments were performed using females and males of Drosophila melanogaster, between 7 and 14 days old. Flies were raised on Iberia food, with 12:12 light-dark cycle, at 25°C. For the CsChR optogenetics experiments, the progeny was reared on Iberia food containing 0.5 mM of all-trans-retinal (ATR) (ApexBio), and adults were collected 2 days after eclosion and reared in Iberia food with 1 mM ATR. Fly lines were sourced from Bloomington Drosophila Stock Center (BDSC), unless differently stated (see table S1). Central complex lines were selected based on previous studies and based on expression reported in Janelia FlyLight project database. PFNd (SS00078), PFNv (SS52628), EPG (SS00090), and PFR (SS54549) lines were reported by Wolff et al. (*41*). Empty Split-Gal4 line (referred to as Control 2) was created by Hampel et al (*43*). To create the hΔB-Split Gal4 line, VT055827-GAL4.DBD and R72B05-p65.AD were recombined, as reported by Lu et al 2022 (*38*). Driver lines for the Mushroom bodies were selected based on expression from Aso et al. (*34*).

Behavior

All experiments were conducted in custom-built arenas. The 50 bl arena was made using three layers of 3D printed PLA parts sandwiched in between two custom-designed PCBs, which served as floor and ceiling. The PCBs were used to deliver the electric shock. We used arenas of 20-120mm diameter, 5mm width, and 5mm height. The arenas were back-illuminated with upward-directed 850 nm LEDs, allowing the top-mounted camera (Allied vision Pike F-1100B) fitted with a 16mm/F1.4 lens (67714 Edmund optics) to record the fly movements at 30 frames per second. LED light source (Thorlabs M625L4, 625 nm and ~20mW $cm^{-2}$ optical power for CsChrimson, 530 nm for GtACR1, Thorlabs M530L4), collimated with an Ø50 mm aspheric condenser lens (Thorlabs COP1-A/B), were used for optogenetic stimulations. The apparatus sits in a temperature-controlled incubator set at 25°C. Arenas were cleaned daily with 70% industrial methylated spirit (Fisher Scientific).

The LabVIEW virtual environment recorded the flies' positions and relayed electric shock and optogenetic stimulations. Previous works describe the tracking and shock-delivery methods (*44,45*). Optogenetic stimulus delivery was controlled by delivering Labview-controlled TTL pulses to T-Cube™ LED Drivers (Thorlabs LEDD1B). We aspirated a single fly into each chamber at the start of each experiment. The flies were given 150-500 s, depending on the size of the chamber, to explore the whole chamber. They were then subjected to an operant training period during which the fly would receive an electric shock (0.4 s long shock every 4.5 s), if they entered a pre-defined part of the chamber. The operant training protocol was adjusted for each chamber size to give the flies sufficient time to escape the shock-associated zone.

Quantification and statistical analyses

We analyzed all data using Matlab R2021b. GraphPad Prism 10 was used for statistical analyses. Full details of statistical analyses can be found in table S2.

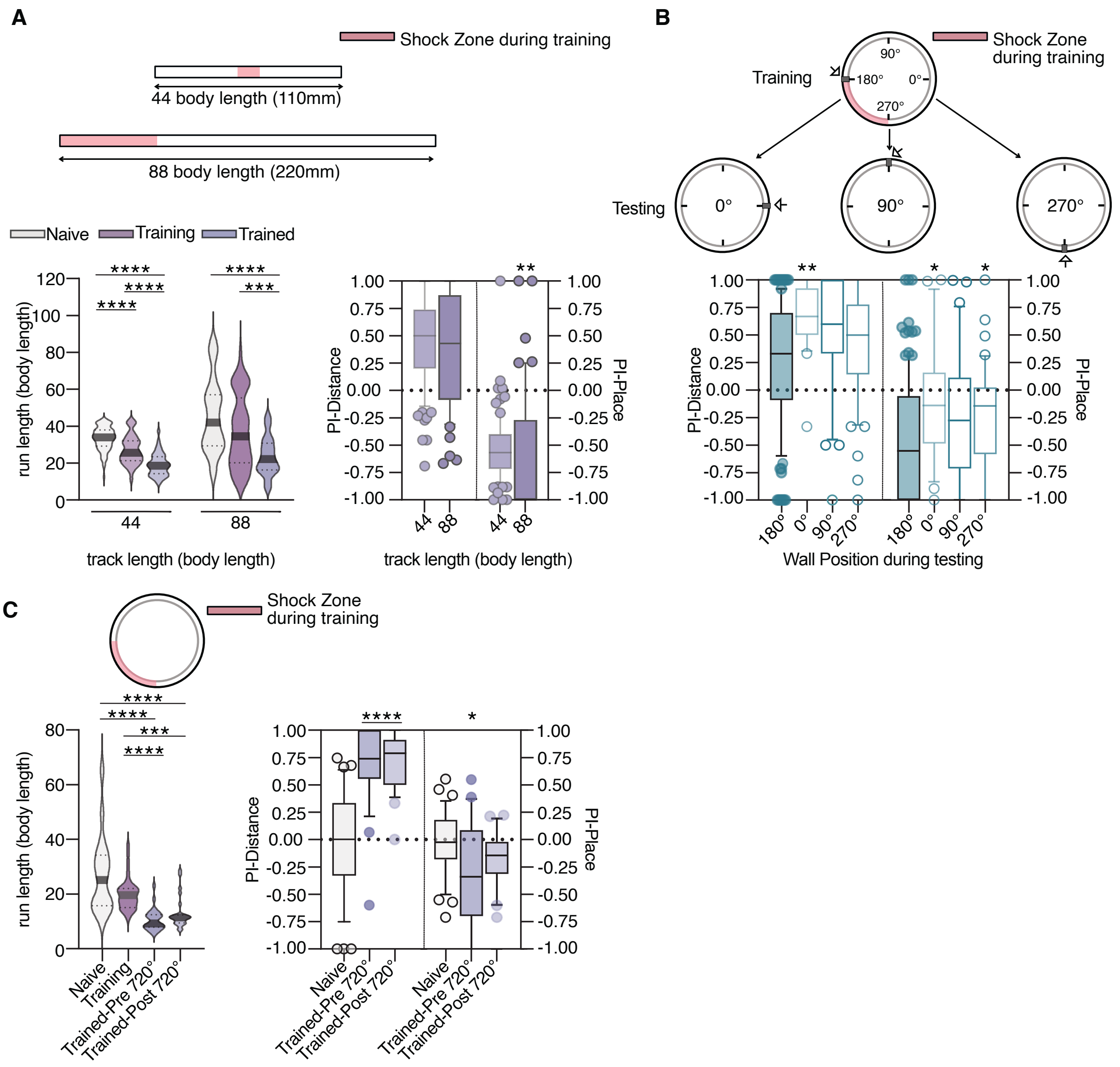


**Fig. S1. Flies can learn in different arena configurations.** (**A**) (Top) Linear arenas of 44 and 88 body lengths; shock zone in pink. (Bottom Left) Flies walked shorter distances between successive turns after training, as in Fig 1B-C. (Bottom Right) Flies showed comparable distance learning in both chamber sizes. (**B**) Moving the partitioning wall (arrowhead) of the 106 body length long annular arena after training to 0°, 90°, or 270° positions. PI- Distance and Place were compared between the static wall (at 180° position) and displaced wall conditions. Place memory was affected by the wall movement. Static wall data from Fig 1E-F is used for the comparison. (**C**) Flies trained in a wall-les annular arena (50 body length long track) showed similar learning to those in Fig.1. Place avoidance dropped after flies made two full rotations along the track.

In all box and whiskers plots, the box extends from 25-75th percentiles with a median line, and whiskers extend from 10-90% percentiles. Data outside these ranges are displayed as individual points. Asterisks indicate significant differences: * $p<0.05$, ** $p<0.01$, *** $p<0.001$, **** $p<0.0001$.

**Table S1. Sources of fly lines used in this study.**

| Resource | Source | Identifier |
|---|---|---|
| *D. melanogaster* UAS-CsChrimson::Venus | Gift from G. Miesenboeck | N/A |
| *D. melanogaster* UAS-GtACR1 w[*]; P{y[+t7.7] w[+mC]=UAS-GtACR1.d.EYFP}attP2 | BDSC | RRID:BDSC_92983 |
| *D. melanogaster* Canton S | Gift from H. Weavers | N/A |
| *D. melanogaster* w[*]; TI{w[+mW.hs]=TI}Orco[1] | BDSC | RRID:BDSC_23129 |
| *D. melanogaster* Empty Split-Gal4w[1118]; P{y[+t7.7] w[+mC]=p65.AD.Uw}attP40; P{y[+t7.7] w[+mC]=GAL4.DBD.Uw}attP2 | BDSC | RRID:BDSC_79603 |
| *D. melanogaster* Empty-Gal4 w[1118]; P{y[+t7.7] w[+mC]=GAL4.1Uw}attP2 | BDSC | RRID:BDSC_68384 |
| *D. melanogaster* PFNd w[1118]; P{y[+t7.7] w[+mC]=R16D01-p65.AD}attP40; P{y[+t7.7] w[+mC]=R15E01-GAL4.DBD}attP2 | BDSC | RRID:BDSC_75854 |
| *D. melanogaster* PFNv w[1118]; P{y[+t7.7] w[+mC]=R22G07-p65.AD}JK73A; P{y[+t7.7] w[+mC]=VT063307-GAL4.DBD}attP2/TM3, Sb[1] | BDSC | RRID:BDSC_76008 |
| *D. melanogaster* hΔB w[1118]; P{y[+t7.7] w[+mC]=R72B05-p65.AD}attP40/CyO; P{y[+t7.7] w[+mC]=VT055827-GAL4.DBD}attP2/TM6B, Tb | (*5*) | N/A |
| *D. melanogaster* w[1118]; P{y[+t7.7] w[+mC]=VT055827-GAL4.DBD}attP2 | BDSC | RRID:BDSC_71851 |
| *D. melanogaster* w[1118]; P{y[+t7.7] w[+mC]=R72B05-p65.AD}attP40; MKRS/TM6B, Tb[1] | BDSC | RRID:BDSC_70939 |
| *D. melanogaster* PFR w[1118]; P{y[+t7.7] w[+mC]=R38B06-p65.AD}attP40; P{y[+t7.7] w[+mC]=VT027015-GAL4.DBD}attP2 | BDSC | RRID:BDSC_86603 |
| *D. melanogaster* EPG w[1118]; P{y[+t7.7] w[+mC]=R19G02-p65.AD}attP40; P{y[+t7.7] w[+mC]=R15C03-GAL4.DBD}attP2 | BDSC | RRID:BDSC_ 75849 |
| *D. melanogaster* KC γd w[1118]; P{y[+t7.7] w[+mC]=R26E07-p65.AD}attP40/CyO; P{y[+t7.7] w[+mC]=R39A11-GAL4.DBD}attP2 | BDSC | RRID:BDSC_ 68323 |
| *D. melanogaster* KC γ w[1118]; P{y[+t7.7] w[+mC]=R19B03-p65.AD}attP40; P{y[+t7.7] w[+mC]=R39A11-GAL4.DBD}attP2 | BDSC | RRID:BDSC_ 68256 |
| *D. melanogaster* KC αβ c w[1118]; P{y[+t7.7] w[+mC]=R13F02-p65.AD}attP40; P{y[+t7.7] w[+mC]=R58F02-GAL4.DBD}attP2 | BDSC | RRID:BDSC_ 68255 |
| *D. melanogaster* KC αβ w[1118]; P{y[+t7.7] w[+mC]=R13F02-p65.AD}attP40; P{y[+t7.7] w[+mC]=R44E04-GAL4.DBD}attP2 | BDSC | RRID:BDSC_ 68291 |

| | | |
|---|---|---|
| *D. melanogaster* KC α’β’ w[1118]; P{y[+t7.7] w[+mC]=R35B12-p65.AD}attP40; P{y[+t7.7] w[+mC]=R26E07-GAL4.DBD}attP2 | BDSC | RRID:BDSC_ 68327 |
| *D. melanogaster* KC αβ s w[1118]; P{y[+t7.7] w[+mC]=R44E04-p65.AD}attP40; P{y[+t7.7] w[+mC]=R26E07-GAL4.DBD}attP2 | BDSC | RRID:BDSC_ 68328 |
| *D. melanogaster* KC αβ p w[1118]; P{y[+t7.7] w[+mC]=R13F02-p65.AD}attP40; P{y[+t7.7] w[+mC]=R85D07-GAL4.DBD}attP2 | BDSC | RRID:BDSC_ 68383 |

**Table S2. Statistical analyses.**

| Figure | Statistical test | Pairwise comparison | Test statistic | *P* | n1 | n2 |
|---|---|---|---|---|---|---|
| **1B** | Ordinary one-way ANOVA | 25bl | F=20.02 | <0.0001 | 61 | 61 |
| | Tukey's test | Naïve vs Training | | 0.3022 | | |
| | Tukey's test | Naïve vs Trained | | <0.0001 | | |
| | Tukey's test | Training vs Trained | | <0.0001 | | |
| | Ordinary one-way ANOVA | 50bl | F=19.28 | <0.0001 | 38 | 38 |
| | Tukey's test | Naïve vs Training | | 0.7781 | | |
| | Tukey's test | Naïve vs Trained | | <0.0001 | | |
| | Tukey's test | Training vs Trained | | <0.0001 | | |
| | Kruskal-Wallis ANOVA | 106bl | H=27.32 | <0.0001 | 124 | 124 |
| | Dunn's test | Naïve vs Training | | >0.99 | | |
| | Dunn's test | Naïve vs Trained | | <0.0001 | | |
| | Dunn's test | Training vs Trained | | <0.0001 | | |
| | Kruskal-Wallis ANOVA | 150bl | H=15.44 | <0.0001 | 19 | 19 |
| | Dunn's test | Naïve vs Training | | 0.0003 | | |
| | Dunn's test | Naïve vs Trained | | 0.1484 | | |
| | Dunn's test | Training vs Trained | | 0.1484 | | |
| **1C** | Linear fit | | Yintercept=7.93 | | | |
| | | | Slope=0.4718 | | | |
| | | | Adj $R^2$=0.3909 | | | |
| **1D** | Kruskal-Wallis ANOVA | | H=3.62 | 0.3053 | | |
| | Dunn's test | 25bl vs. 50bl | Z=1.722 | 0.5106 | 64 | 38 |
| | | 25mm vs. 106bl | Z=0.4062 | >0.9999 | 64 | 124 |
| | | 25bl vs. 150bl | Z=0.214 | >0.9999 | 64 | 19 |
| | | 50bl vs. 106bl | Z=1.565 | 0.706 | 38 | 124 |
| | | 50bl vs. 150bl | Z=1.454 | 0.8756 | 38 | 19 |
| | | 106bl vs. 150bl | Z=0.4807 | >0.9999 | 124 | 19 |
| **1E; left** | Brown-Forsythe ANOVA | Naïve | F=5.988 (3.000, 102.9) | 0.0008 | | |
| | Dunnett's test | 25bl vs. 50bl | t=0.2045 | >0.9999 | 64 | 38 |
| | Dunnett's test | 25mm vs. 106bl | t=3.411 | 0.0049 | 64 | 124 |

| | | | | | | |
|---|---|---|---|---|---|---|
| | Dunnett's test | 25bl vs. 150bl | t=2.337 | 0.147 | 64 | 19 |
| | Dunnett's test | 50bl vs. 106bl | t=3.062 | 0.0168 | 38 | 124 |
| | Dunnett's test | 50bl vs. 150bl | t=2.18 | 0.1989 | 38 | 19 |
| | Dunnett's test | 106bl vs. 150bl | t=0.3804 | 0.9992 | 124 | 19 |
| **1E; right** | Kruskal-Wallis ANOVA | Trained | H=16.82 | 0.0008 | | |
| | Dunn's test | 25bl vs. 50bl | Z=0.7042 | >0.9999 | 64 | 38 |
| | Dunn's test | 25bl vs. 106bl | Z=0.9081 | >0.9999 | 64 | 123 |
| | Dunn's test | 25bl vs. 150bl | Z=3.261 | 0.0067 | 64 | 19 |
| | Dunn's test | 50bl vs. 106bl | Z=0.02298 | >0.9999 | 38 | 123 |
| | Dunn's test | 50bl vs. 150bl | Z=3.545 | 0.0024 | 38 | 19 |
| | Dunn's test | 106bl vs. 150bl | Z=4.024 | 0.0003 | 123 | 19 |
| **1F; left** | Kruskal-Wallis ANOVA | Place | H=41.31 | <0.0001 | | |
| | Uncorrected Dunn's | Naive vs. 0 | Z=2.635 | 0.0084 | 48 | 48 |
| | Uncorrected Dunn's | Naive vs. 10 | Z=2.48 | 0.0132 | 48 | 48 |
| | Uncorrected Dunn's | Naive vs. 20 | Z=3.566 | 0.0004 | 48 | 47 |
| | Uncorrected Dunn's | Naive vs. 30 | Z=4.47 | <0.0001 | 48 | 48 |
| | Uncorrected Dunn's | Naive vs. 40 | Z=5.026 | <0.0001 | 48 | 47 |
| | Uncorrected Dunn's | Naive vs. 50 | Z=4.033 | <0.0001 | 48 | 45 |
| | Uncorrected Dunn's | Naive vs. 55 | Z=5.196 | <0.0001 | 48 | 44 |
| **1F; right** | Ordinary one-way ANOVA | Place | F=2.331 | 0.0244 | | |
| | Uncorrected Fisher's | Naive vs. 0 | t=2.767 | 0.0059 | 48 | 48 |
| | Uncorrected Fisher's | Naive vs. 10 | t=0.5931 | 0.5535 | 48 | 48 |
| | Uncorrected Fisher's | Naive vs. 20 | t=1.03 | 0.3036 | 48 | 47 |
| | Uncorrected Fisher's | Naive vs. 30 | t=0.2987 | 0.7654 | 48 | 48 |
| | Uncorrected Fisher's | Naive vs. 40 | t=0.2581 | 0.7965 | 48 | 47 |
| | Uncorrected Fisher's | Naive vs. 50 | t=0.5428 | 0.5876 | 48 | 45 |
| | Uncorrected Fisher's | Naive vs. 55 | t=0.09995 | 0.9204 | 48 | 44 |

| | | | | | | |
|---|---|---|---|---|---|---|
| **1G** | Unpaired t-test | Distance: orco1 vs CS | t=0.8753, df=93 | 0.3837 | 49 | 46 |
| | Mann-Whitney test | Place: orco1 vs CS | U= 642 | 0.0002 | 49 | 46 |
| **2A** | Kruskal-Wallis ANOVA | Distance | H=10.96 | 0.3606 | | |
| | Dunn's test | Control1 vs. control2 | Z=0.2246 | >0.9999 | 24 | 70 |
| | Dunn's test | Control1 vs. $\alpha\beta_c$ | Z=0.3409 | >0.9999 | 24 | 42 |
| | Dunn's test | Control1 vs. γ | Z=0.8673 | >0.9999 | 24 | 35 |
| | Dunn's test | Control1 vs. αβ | Z=0.9156 | >0.9999 | 24 | 44 |
| | Dunn's test | Control1 vs. γd | Z=0.6823 | >0.9999 | 24 | 45 |
| | Dunn's test | Control1 vs. α'β' | Z=0.3905 | >0.9999 | 24 | 37 |
| | Dunn's test | Control1 vs. $\alpha\beta_s$ | Z=0.774 | >0.9999 | 24 | 47 |
| | Dunn's test | Control1 vs. $\alpha\beta_p$ | Z=0.556 | >0.9999 | 24 | 39 |
| | Ordinary one-way ANOVA | Place | F=1.324 | 0.2149 | | |
| | Dunnett's test | Control1 vs. control2 | q=0.2755 | >0.9999 | 69 | 24 |
| | Dunnett's test | Control1 vs. $\alpha\beta_c$ | q=0.6123 | 0.9989 | 69 | 42 |
| | Dunnett's test | Control1 vs. γ | q=0.1004 | >0.9999 | 69 | 35 |
| | Dunnett's test | Control1 vs. αβ | q=1.786 | 0.4652 | 69 | 44 |
| | Dunnett's test | Control1 vs. γd | q=0.7246 | 0.9958 | 69 | 45 |
| | Dunnett's test | Control1 vs. α'β' | q=1.368 | 0.781 | 69 | 37 |
| | Dunnett's test | Control1 vs. $\alpha\beta_s$ | q=2.376 | 0.1433 | 69 | 47 |
| | Dunnett's test | Control1 vs. $\alpha\beta_p$ | q=0.3899 | >0.9999 | 69 | 39 |
| **2B** | Kruskal-Wallis ANOVA | Distance | H=19.31 | 0.0017 | | |
| | Dunn's test | Control vs. PFNd | Z=2.833 | 0.0231 | 117 | 79 |
| | Dunn's test | Control vs. PFNv | Z=3.682 | 0.0012 | 117 | 120 |
| | Dunn's test | Control vs. EPG | Z=1.497 | 0.6725 | 117 | 99 |
| | Dunn's test | Control vs. hΔB | Z=2.954 | 0.0157 | 117 | 56 |
| | Dunn's test | Control vs. PFR | Z=1.258 | >0.9999 | 117 | 105 |
| | Kruskal-Wallis ANOVA | Place | H=17.5 | 0.0036 | | |
| | Dunn's test | Control vs. PFNd | Z=2.24 | 0.1255 | 117 | 79 |
| | Dunn's test | Control vs. PFNv | Z=3.661 | 0.0013 | 117 | 120 |
| | Dunn's test | Control vs. EPG | Z=0.7308 | >0.9999 | 117 | 99 |
| | Dunn's test | Control vs. hΔB | Z=1.873 | 0.3053 | 117 | 56 |
| | Dunn's test | Control vs. PFR | Z=0.8001 | >0.9999 | 117 | 105 |
| **2C** | Kruskal-Wallis ANOVA | Distance | H=5.647 | 0.3421 | | |
| | Dunn's test | Control vs. PFNd | Z=0.02743 | >0.9999 | 109 | 59 |
| | Dunn's test | Control vs. PFNv | Z=1.383 | 0.8334 | 109 | 85 |

| | | | | | | |
|---|---|---|---|---|---|---|
| | Dunn's test | Control vs. EPG | Z=0.02743 | >0.9999 | 109 | 59 |
| | Dunn's test | Control vs. hΔB | Z=0.07036 | >0.9999 | 109 | 61 |
| | Dunn's test | Control vs. PFR | Z=1.173 | >0.9999 | 109 | 73 |
| | Kruskal-Wallis ANOVA | Place | H=2.27 | 0.8107 | | |
| | Dunn's test | Control vs. PFNd | Z=0.6148 | >0.9999 | 109 | 59 |
| | Dunn's test | Control vs. PFNv | Z=0.5906 | >0.9999 | 109 | 85 |
| | Dunn's test | Control vs. EPG | Z=0.6148 | >0.9999 | 109 | 59 |
| | Dunn's test | Control vs. hΔB | Z=0.4891 | >0.9999 | 109 | 61 |
| | Dunn's test | Control vs. PFR | Z=0.2871 | >0.9999 | 109 | 73 |
| **3A** | Mann-Whitney test | Distance hΔB vs. Control | U=911 | <0.0001 | 59 | 58 |
| **3C** | Mann-Whitney test | Distance hΔB vs. Control | U=1055 | 0.0040 | 57 | 54 |
| | Unpaired t-test | Distance hΔB vs. Control | t=0.7015, df=204 | 0.4838 | 91 | 115 |
| **3D** | Mann-Whitney test | Distance hΔB vs. Control | U=1783 | 0.9821 | 55 | 65 |
| | Mann-Whitney test | Distance hΔB vs. Control | U=1594 | 0.0087 | 93 | 47 |
| | Mann-Whitney test | Distance hΔB vs. Control | U=1565 | 0.2830 | 57 | 62 |
| **4C** | Kruskal-Wallis ANOVA | | H=70.18 | <0.0001 | | |
| | Dunn's test | Start: Control vs. hΔB | Z=3.893 | 0.0003 | 146 | 152 |
| | Dunn's test | Middle: Control vs. hΔB | Z=6.620 | <0.0001 | 481 | 289 |
| | Dunn's test | End: Control vs. hΔB | Z=3.223 | 0.0038 | 164 | 169 |
| **4D** | Kruskal-Wallis ANOVA | | H=716.8 | <0.0001 | | |
| | Dunn's test | Start: Control vs. hΔB | Z=2.475 | 0.0400 | 158 | 231 |
| | Dunn's test | Middle: Control vs. hΔB | Z=8.171 | <0.0001 | 434 | 466 |
| | Dunn's test | End: Control vs. hΔB | Z=2.200 | 0.0833 | 166 | 241 |
| **S1A; left** | Kruskal-Wallis ANOVA | Distance | H=115.8 | <0.0001 | | |
| | Dunn's test | Naïve vs Training | Z=4.799 | <0.0001 | 89 | 89 |
| | Dunn's test | Naïve vs Trained | Z=10.74 | <0.0001 | 89 | 89 |
| | Dunn's test | Training vs Trained | Z=5.942 | <0.0001 | 89 | 89 |

| | | | | | | |
|---|---|---|---|---|---|---|
| | Kruskal-Wallis ANOVA | Distance | H=39.11 | <0.0001 | | |
| | Dunn's test | Naïve vs Training | Z=2.340 | 0.0579 | 58 | 58 |
| | Dunn's test | Naïve vs Trained | Z=6.192 | <0.0001 | 58 | 58 |
| | Dunn's test | Training vs Trained | Z=3.852 | 0.0004 | 58 | 58 |
| **S1A; right** | Unpaired t-test | Distance 88bl vs 44bl | t=0.9646, df=139 | 0.3364 | 52 | 89 |
| | Mann-Whitney test | Place 88bl vs 44bl | U=1586 | 0.0045 | 50 | 89 |
| **S1B** | Kruskal-Wallis ANOVA | Distance | H=12.88 | 0.0049 | | |
| | Dunn's test | Static vs. 270° | Z=1.290 | 0.5910 | 123 | 51 |
| | Dunn's test | Static vs. 90° | Z=2.039 | 0.1242 | 123 | 31 |
| | Dunn's test | Static vs. 0° | Z=3.241 | 0.0036 | 123 | 21 |
| | Kruskal-Wallis ANOVA | Place | H=13.44 | 0.0038 | | |
| | Dunn's test | Static vs. 270° | Z=2.793 | 0.0157 | 123 | 49 |
| | Dunn's test | Static vs. 90° | Z=1.704 | 0.2650 | 123 | 31 |
| | Dunn's test | Static vs. 0° | Z=2.764 | 0.0171 | 123 | 21 |
| **S1C; left** | Kruskal-Wallis ANOVA | | H=59.56 | <0.0001 | | |
| | Dunn's test | Naive vs. Training | Z=1.192 | >0.9999 | 35 | 35 |
| | Dunn's test | Naive vs. Trained -Pre720º | Z=6.549 | <0.0001 | 35 | 26 |
| | Dunn's test | Naive vs. Trained-Post720º | Z=5.164 | <0.0001 | 35 | 24 |
| | Dunn's test | Training vs. Trained-Pre720º | Z=5.445 | <0.0001 | 35 | 26 |
| | Dunn's test | Training vs. Trained-Post720º | Z=4.088 | 0.0003 | 35 | 24 |
| | Dunn's test | Trained-Pre720º vs. Trained-Post720º | Z=1.153 | >0.9999 | 26 | 24 |
| **S1C; right** | Kruskal-Wallis ANOVA | | H=40.02 | <0.0001 | | |
| | Dunn's test | Naive vs. Trained -Pre720º | Z=5.393 | <0.0001 | 35 | 26 |
| | Dunn's test | Naive vs. Trained-Post720º | Z=5.386 | <0.0001 | 35 | 24 |
| | Ordinary one-way ANOVA | | F=4.230 | 0.0178 | | |
| | Dunnett's test | Naive vs. Trained -Pre720º | q=2.843 | 0.0109 | 35 | 27 |

| | | | | | | |
|---|---|---|---|---|---|---|
| | Dunnett's test | Naive vs. Trained-Post720º | q=1.754 | 0.1505 | 35 | 24 |
